# Multi-omic latent variable data integration reveals multicellular structure pathways associated with resistance to tuberculin skin test (TST)/interferon gamma release assay (IGRA) conversion in Uganda

**DOI:** 10.1101/2024.11.05.622020

**Authors:** Madison S Cox, Kimberly A Dill-McFarland, Jason D Simmons, Penelope Benchek, Harriet Mayanja-Kizza, W Henry Boom, Catherine M Stein, Thomas R Hawn

## Abstract

Understanding the mechanisms of early clearance of *Mycobacterium tuberculosis* (Mtb) may illuminate new therapeutic strategies for tuberculosis (TB). We previously found genetic, epigenetic, and transcriptomic signatures associated with resistance (resister, RSTR) to tuberculin skin test (TST)/interferon gamma release assay (IGRA) conversion among highly exposed TB contacts. We hypothesized that integration of these datasets with multi-omic latent factor methods would detect pathways differentiating RSTR patients from those with latent infection (LTBI) which were not differentiated by individual dataset analyses. We pre-filtered and scaled features with the largest change between LTBI and RSTR groups for 126 patients with data in at least two of five data modalities: single nucleotide polymorphisms (SNP), monocyte RNAseq (baseline and Mtb-stimulated conditions), and monocyte epigenetics (methylation and ATAC-seq). Using multiomic latent factor analysis (MOFA), we generated ten latent factors on the subset of 33 patients with all five datasets available, four of which were different between LTBI and RSTR (FDR < 0.1). Factor 4, which was the most integrated of the significant factors, showed the greatest difference between RSTR and LTBI groups (FDR < 0.001). Three additional latent factor data integration methods also distinguished the RSTR and LTBI groups and identified overlapping features with MOFA. Using pathway analysis and a cluster-based enrichment method, we identified biologic functions associated with latent factors and found that MOFA Factors 2-4 include functions related to cell-cell adhesion, cell shape, and development of multicellular structures. In summary, latent variable integration methods uncovered signatures associated with resistance to TST/IGRA conversion that were not detected within individual dataset analyses and included pathways associated with cellular interactions and multicellular structures.

**Author Summary:** Tuberculosis (TB) is a leading cause of preventable death worldwide. Previous research has identified some genetic and molecular patterns that are linked to resistance to TB in people who are highly exposed to the bacterium that causes the disease but do not develop an infection. We took genetic, epigenetic, and gene expression data from people who were not infected after exposure to TB within in their household and compared them with people who did develop a latent form of TB disease after similar exposure. We used a special method to analyze these datasets in an integrated way, rather than separately. This approach revealed four key factors that clearly distinguished between the resistant individuals and those with latent infection. One of these factors, in particular, showed a strong difference. We found that these factors were related to important biological processes such as how cells stick together, their shape, and how they form tissues. This method allowed us to see new patterns in the biological pathways linked to TB resistance that were not evident when looking at each type of data on its own.

## Introduction

Tuberculosis (TB) is a leading cause of global mortality including 1.3 million deaths among 10.6 million cases reported in 2022(1). Following heavy exposure to *Mycobacterium tuberculosis* (Mtb), a range of outcomes occurs including TB disease, asymptomatic or latent TB infection (LTBI) defined clinically as a positive tuberculin skin test (TST) or IFNγ release assay (IGRA), and resistance to TST/IGRA conversion (RSTR) that may represent clearance of infection through IFNγ-independent mechanisms(2–4).

Immunologic and genetic mechanisms of resistance to Mtb infection following close contact have been investigated in several cohorts(3,5–8) including a long-term household contacts study in Uganda(2,4), a country with a high incidence of Mtb infection(9). Genetic(10,11), epigenetic(12), and transcriptional(13,14) signatures differentiating LTBI and RSTR subjects have been described in monocyte-derived data from this cohort, pointing to several possible mechanisms of resistance within the inflammatory response including TNF responses(11,13) and lipid metabolism(12,14). However, there is little agreement across these data modalities in terms of specific genes or pathways that might be investigated as potential therapeutic targets. An integrated analysis of these datasets may help to further identify pathways and features that differentiate LTBI and RSTR subjects, generating new lines of inquiry for investigation into the mechanisms underlying resistance to Mtb infection(15,16).

In this work, we utilize several multi-omic latent variable integration methods to identify driving sources of variation across data modalities. The primary integration method used is MOFA+, an unsupervised factor analysis method(17). MOFA+ and similar integrative computational methods can provide mechanistic insights above and beyond traditional -omic data analyses by revealing functional pathways whose regulation may span across more than one step in the cascade from chromosome to protein or metabolite. These methods have aided in novel biomarker identification, classification of disease subtypes, and discovery of candidate drug targets in various diseases with complex mechanisms(18,19). By integrating genetics with monocyte-derived methylation, chromatin accessibility, and transcriptomic datasets from the Uganda resister cohort, we explored mechanisms of resistance to TST/IGRA conversion that were not detected in the previous analyses of each independent dataset.

## Methods

### Cohort

Patients with culture-positive pulmonary tuberculosis (TB) were recruited as part of the Kawempe Community Health Study from 2002 to 2012 in Kampala, Uganda(4). Protocols were approved by University Hospitals Institutional Review Board (IRB) at University Hospitals Cleveland Medical Center (IRB number 10-01-25) and the National HIV/AIDS Research Committee and Uganda National Council for Science and Technology in Uganda (reference number MV 658). Household contacts of index TB cases were then enrolled and followed for 2 years by tuberculin skin tests (TST). A subset of TST-negative and matched TST-positive individuals were retraced from 2014 and 2017 and re-assessed by TST as well as IFNγ release assays (IGRA) for an additional 2 years(2). Latent tuberculosis infection (LTBI) was defined as individuals with fully concordant positive TST and IGRA tests. Resisters (RSTR) were defined as concordant negative TST and IGRA. All participants were at least 15 years old at the time of retracing, HIV-negative, and gave written, informed consent, approved by the institutional review boards of their associated institutions. Previously generated data include genetic association studies with single nucleotide polymorphisms (SNPs)(11), as well as chromatin accessibility (ATAC-seq)(12), methylation(12), and transcriptional responses (RNA-seq)(13,14) in monocytes.

### Kinship

Genotypes were determined using the Illumina MEGA^EX^ array containing 2 million single nucleotide polymorphism (SNP) probes or Infinium OmniExpress BeadChip containing 710,000 probes as previously reported(11). SNPs were filtered for minor allele frequency (MAF > 0.05), call rate (> 0.95), Hardy-Weinberg Equilibrium (P < 1×10^-6^), and linkage disequilibrium (LD R^2^ < 0.1 in 50 bp windows with a 5 bp slide) in PLINK2(20). This yielded 63812 filtered SNPs for kinship calculation. Pairwise kinship was calculated using the robust King method for identity-by-descent (IBD, SNPRelate v1.22.0)(21) and a genetic relationship matrix (GRM, GENESIS v2.18.0(22)).

### Data preprocessing

To examine integrated profiles that distinguish RSTR and LTBI clinical groups with MOFA, we selected five previously published datasets: monocyte RNA-seq(13,14) (media condition and Mtb-stimulated), monocyte methylation(12) (media condition), monocyte ATAC-seq(12) (media condition), and SNPs(11). The study design including data processing and analysis is summarized in Figure 1. Collectively, these SNPs, methylation probes, ATAC-seq peaks, and RNA-seq genes are referred to as features for integration. Features not annotated to a known gene (GRCh38)(23) were omitted. In the methylation data, outliers > 4 SD from the overall mean were rescaled to 4 SD. In the continuous datasets (RNA-seq [media and Mtb-stimulated], ATAC-seq, methylation), log2 fold changes were calculated for RSTR vs. LTBI. For ordinal SNP data, fold changes in minor allele frequencies were calculated by for RSTR/LTBI. For the larger datasets (features > 1×10^5),^ the top 1% of features with the greatest absolute log2 fold change were selected for downstream analysis (SNPs = 6609, methylation = 5349). For the smaller datasets (features < 1×10^5^), the features in the top 10% by greatest absolute log2 fold change were selected (RNA-seq = 1398, ATAC-seq = 2466). Subjects missing any of the five integrated datasets were omitted. This resulted in 33 subjects and 1.7×10^4^ total features for integration. Data completeness for the full cohort of 126 patients is summarized in Supplemental Figure 1.

**Figure 1.**
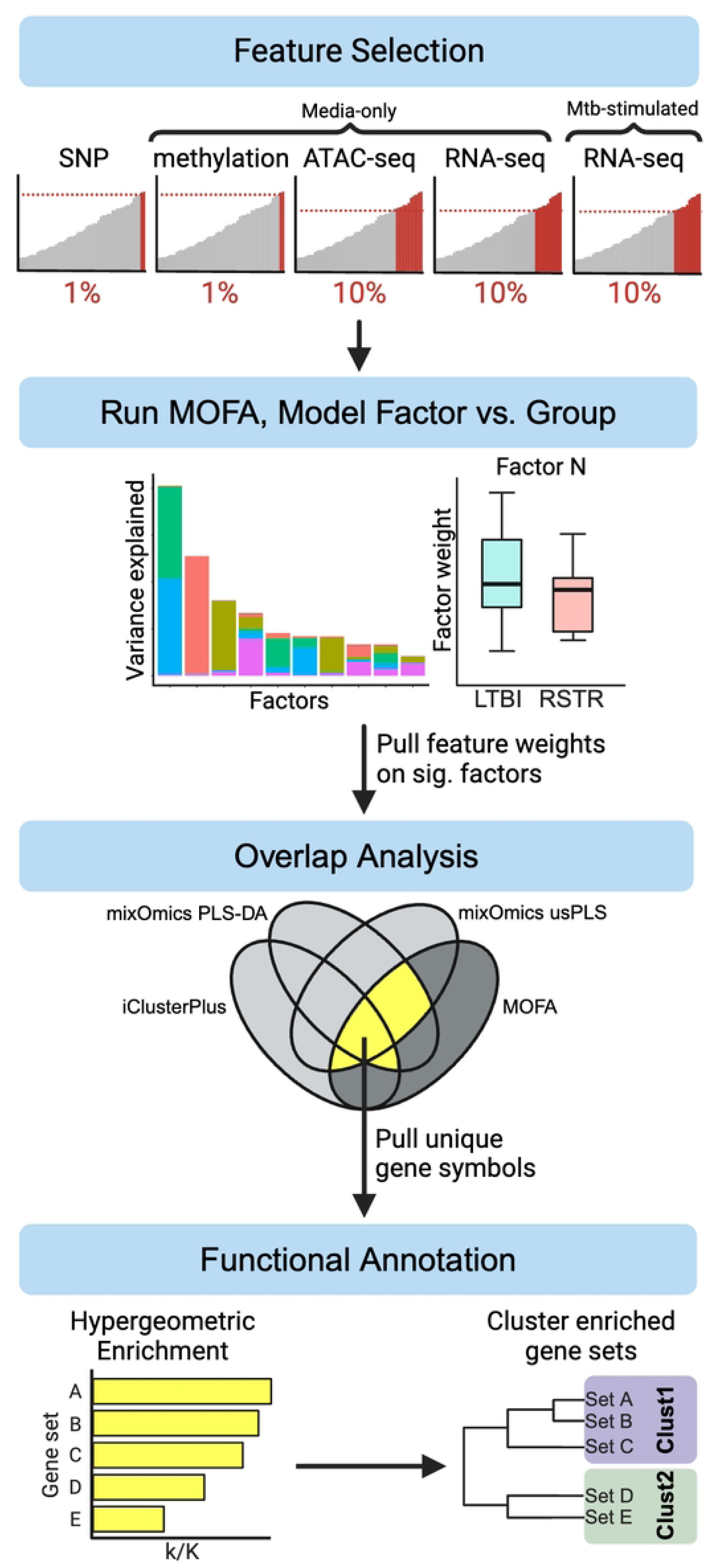
Data processing and analysis workflow for multi-omics integration in the Uganda RSTR cohort. This analysis consisted of four major parts: (1) input feature selection, (2) creation of the MOFA factors, (3) comparison of top-weighted features on significant factors with those from other latent variable integration methods and selection of features highlighted by MOFA and at least two additional methods, (4) and factor annotation based on functional enrichment of those reduced feature lists and summary of enrichment results by gene set clustering.

### MOFA implementation

MOFA infers a set of latent factors to capture sources of variability across and within data modalities with different underlying data structures and distributions(17). Factor generation in MOFA is unsupervised and results in matrices of factor loadings by feature for each of the integrated datasets, as well as factor loadings by subject. From the 5 datasets filtered from 33 subjects, ten latent factors were generated using the MOFA+ package(17) in R v4.2.3(24) using default parameters apart from scale_views being set to TRUE to allow for the different datasets to be scaled to equal variance and the function-internal random seed being set to improve reproducibility. RSTR and LTBI MOFA factor values were compared using a mixed effects models corrected for age, sex, and genetic kinship in kimma(25). Functional annotation was performed on factors that differed between RSTR and LTBI (FDR < 0.2). Data sets that explained

>5% of variance on a factor were considered for functional characterization of that factor. The top five features by MOFA weight in each dataset were subjected to hypergeometric enrichment against MSigDB databases(26) as described below.

### Additional latent variable integration methods

To reduce the feature lists to more specifically characterize functions represented by MOFA factors, an overlap analysis was performed with several alternative multi-omic latent variable integration methods. Three non-MOFA methods were implemented. The first was a multiblock unsupervised partial-least-squares analysis performed using the mixOmics::block.pls function in the canonical mode with Mtb-stimulated RNA-seq designated as the response dataset(27). The design matrix included zeroes on the diagonal and 0.1 on the off-diagonal; two latent variables were generated. Like MOFA, this is an unsupervised method that generates latent factors naïve to sample group. Second, a multiblock partial-least-squares discriminant analysis was performed with the same structure of the design matrix using the mixOmics::block.plsda function to generate two latent variables(27). This is a supervised method that creates factors that best separate LTBI and RSTR groups. Lastly, the unsupervised, graph-based method iClusterPlus was used to generate three latent variables with the function iClusterBayes(28,29). Model tuning was performed using the function tune.iClusterBayes, and the final model was built with default parameters for the Markov Chain Monte Carlo sampling apart from: number of burn-in iterations set to 18,000, number of draws set to 12,000, and prior gamma probability set to 0.5 for all five input datasets.

### Multi-method feature selection

Models were generated with kimma to compare mixOmics and iClusterPlus latent factors by LTBI/RSTR sample group with sex, age, and kinship correction(25). The most extreme 10% of features on each of the significant latent variables (FDR < 0.2) from each of the five datasets were selected to look for overlap with the top MOFA features on RSTR significant factors.

Top MOFA features were selected from datasets explaining at least 5% of variance on factors that significantly differed in RSTR and LTBI. A proportion of the most extreme features were selected equal to (0.25 * *P_d_*), where *P_d_* is the proportion of variance explained by the dataset on that factor. Features were selected from this reduced MOFA feature list that also occurred in the extreme feature list for at least two out of three of the non-MOFA integration methods (Figure 1). The gene annotations associated with these overlapping features were subjected to hypergeometric enrichment using MSigDB as described below.

### Comparison with individual data set analyses

MOFA reduced feature lists for each factor were compared to results from the individual analyses of the integrated datasets. For RNA-seq data, expression was modeled with respect to Mtb stimulation (media-only and Mtb-stimulated) and RSTR status. Features significant for the interaction term (Mtb:RSTR) of that model (FDR < 0.2) were selected for comparison with the MOFA reduced feature lists(13). For the ATAC-seq data, the two peaks that differed (FDR < 0.2) between LTBI and RSTR were used(12). Methylation was assessed both as differentially methylated probes and probes within differentially methylated regions previously defined by DMRcate. For this analysis, methylation features significant under either scheme were included (FDR < 0.2)(12). SNPs from the reduced feature lists were queried from the GWAS dataset and compared with the 40 SNPs that passed the 5×10^-5^ significance threshold for that analysis(11).

### Functional enrichment

Hypergeometric enrichment of protein-coding genes in top-weighted MOFA feature lists and gene lists generated in the overlap analysis were performed using the SEARchways package in R(30). For MOFA Factors 1 and 2, the flexEnrich function was used. For MOFA Factors 3 and 4, there were a number of features with more than one gene annotation, necessitating the use of the iterEnrich function to account for annotation of features to multiple HGNC symbols with random subsampling. Enrichment was tested against the Hallmark(26), C2 curated gene sets (Canonical Processes), and C5 gene ontology gene sets (Biological Processes)(31,32) databases from the Molecular Signatures Database. In all cases, a minimum overlap threshold of three was imposed between the query and pathway gene lists. Additionally, within each analysis the minimum gene set size considered was 10, and the maximum was three standard deviations over the mean gene set size for each database (Hallmark = 386.6, C2 CP = 361.7, C5 GO:BP = 786.2).

In order to summarize pathway enrichment results, significant gene sets (FDR < 0.2) were subjected to hierarchical clustering based on the overlap coefficient(33) calculated on pathway gene membership. Clusters were generated using a tree-cut height of 0.8. Cluster annotation was based on a combination of the largest gene set within each cluster and word cloud diagrams built on member gene set names and descriptions(34).

## Results

To identify new biologic signatures that distinguish RSTR and LTBI phenotypes, we integrated five data sets previously generated from monocytes from household contacts of Mtb cases in Uganda (media-only and Mtb-stimulated RNAseq, ATACseq, methylation, SNPs, Figure 1). Multi-omic factor analysis was applied, creating latent factors that describe axes of heterogeneity that can span across input data modalities. Once these factors have been identified, features with the high absolute weight on the factors can be used to relate them to etiology.

### Dataset selection and preprocessing

The total number of raw features across all five datasets was 1,526,259 with 1,250,370 annotated to known genes. Seven methylation values fell > 4SD from the mean of all methylation data and were rescaled. Data was pre-filtered to the features with the largest difference between LTBI and RSTR groups per dataset for 126 patients with data in at least two of the five data modalities. The final dataset contained 1,398 genes for each of the media-only and Mtb-stimulated RNA-seq datasets, 5,349 methylation probes, 2,466 ATAC-seq peaks, and 6,609 SNPs for a total of 17,220 features across all data types (Table 1). MOFA factors were then generated on the subset of patients with all five datasets available (in Table 2, n = 33).

**Table 1.** Summary of features used in data integration.

| Category | Dataset | Number of features passing dataset QC | Number of features annotated to a known gene | Percent of features selected in preprocessing | Number of features in integrated dataset |
| --- | --- | --- | --- | --- | --- |
| Media-only | RNA-seq | 1.4 x 10<sup>4</sup> | 1.4 x 10<sup>4</sup> | 10% | 1398 |
|  | Methylation | 7.3 x 10<sup>5</sup> | 5.2 x 10<sup>5</sup> | 1% | 5349 |
|  | ATAC-seq | 4.2 x 10<sup>4</sup> | 2.5 x 10<sup>4</sup> | 10% | 2466 |
| Mtb-stimulated | RNA-seq | 1.4 x 10<sup>4</sup> | 1.4 x 10<sup>4</sup> | 10% | 1398 |
| Host-dependent | SNP | 7.3 x 10<sup>5</sup> | 6.7 x 10<sup>5</sup> | 1% | 6609 |
| **Total** |  | **1.5 x 10<sup>6</sup>** | **1.3 x 10<sup>6</sup>** |  | **1.7 x 10<sup>4</sup>** |

**Table 2.** Demographic data and group membership of subjects in input datasets.

|  | LTBI | RSTR | p-value<sup>A</sup> |
| --- | --- | --- | --- |
| N subjects | 15 | 18 |  |
| Median Age at Enrollment (IQR) | 15 (8.5) | 12.5 (7.5) | 0.277 |
| Median Age at Sample Collection (IQR) | 23 (4) | 21.5 (7.5) | 0.293 |
| Sex, % Male (n/N) | 66% (10/15) | 39% (7/18) | 0.215 |
| Median BMI (IQR)<sup>B</sup> | 23.6 (4.0) | 21.2 (6.6) | 0.546 |
| Median Exposure Score at Enrollment (IQR) | 6 (1.5) | 6 (1.0) | 0.690 |
| BCG scar, % Yes (n/N)<sup>B</sup> | 60% (9/15) | 61% (11/18) | 0.981 |
| % HIV+ | 0 | 0 |  |
| Relatedness Within Phenotype |  |  |  |
| Mean 3° or Closer Per Person (SD) | 0.53 (0.52) | 0.33 (0.49) | 0.263 |
| Mean 1° or Closer Per Person (SD) | 0.13 (0.35) | 0 (0) | 0.126 |

<sup>A</sup> P-values calculated by Wilcoxon rank-sum tests for continuous variables and Pearson's chi-squared tests for categorical variables.
<sup>B</sup> NA values were present in these variables. For BMI, the one NA value was in the RSTR group and was excluded from summary statistics. In % BCG scar, the 9 NA values across groups (RSTR = 5, LTBI = 4) were treated as a third group in addition to "Yes" and "No" for the statistical test.

### MOFA latent factors distinguish RSTR and LTBI groups

Features from each dataset were pre-filtered to those with the greatest fold-change difference between LTBI and RSTR. Using this filtered feature list, we generated ten latent factors in MOFA (Supplemental Figures 2 & 3), four of which were different between LTBI and RSTR (Figure 2, Supplemental Table 1, FDR < 0.1). Variance on Factor 1 was primarily explained by features from the two RNA-seq datasets, Factor 2 from the ATAC-seq dataset, and Factor 3 from the methylation and SNP datasets. In contrast, Factor 4 better integrated the input datasets, with > 5% variance explained by each of the SNP, methylation, Mtb-stimulated RNA-seq, and ATAC-seq datasets. Strikingly, Factor 4 showed near perfect discrimination between RSTR and LTBI groups, with the LTBI group associated with negative values on the factor and the RSTR group with positive values on the factor (FDR < 0.001). Together, these analyses suggest that MOFA identified factors which differentiate the RSTR and LTBI groups with features from multiple datasets (Supplemental Figures 4 & 5).

**Figure 2.**
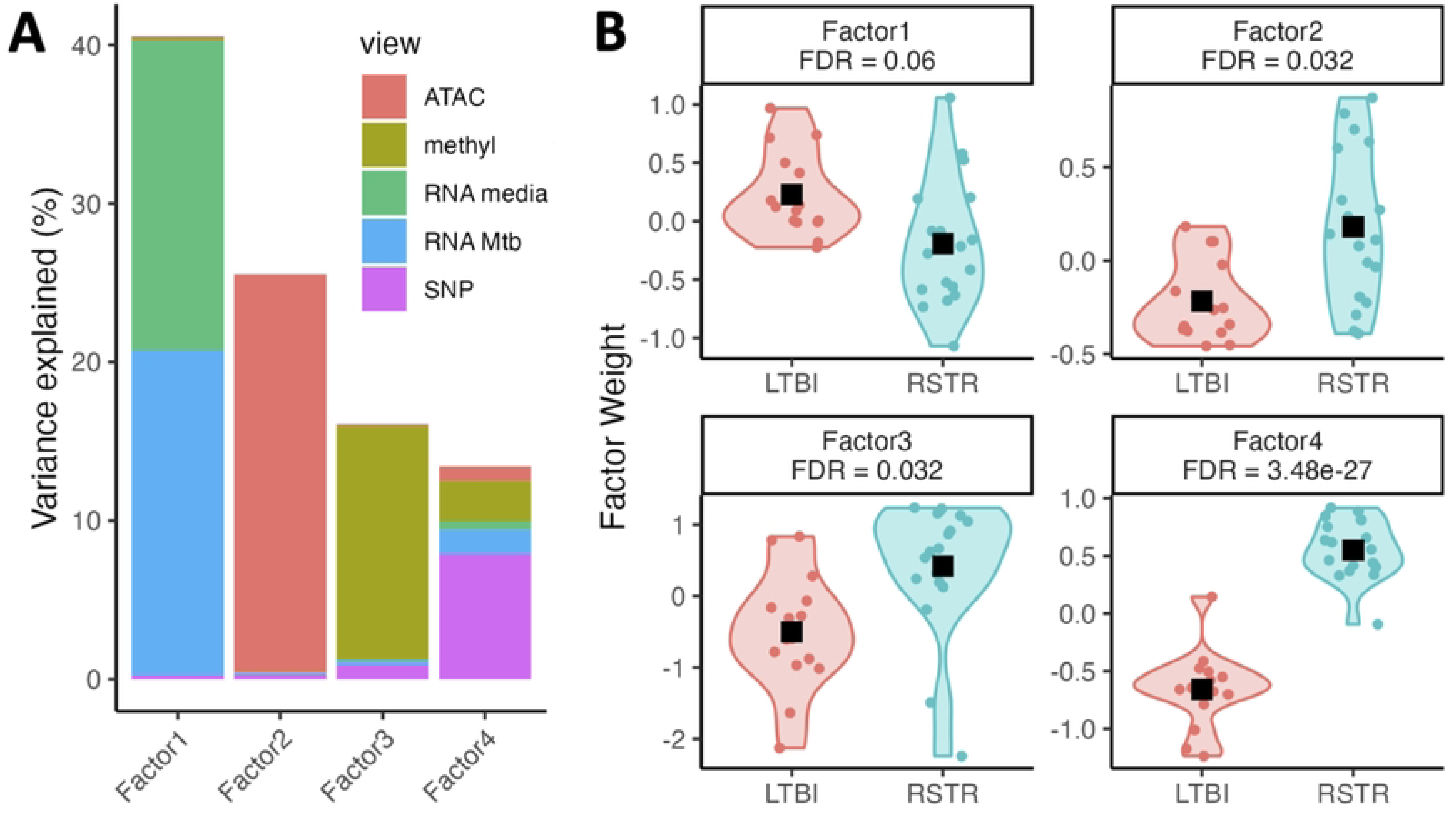
Four MOFA factors differ between RSTR and LTBI. (A) The variance explained by each dataset on each of the first four latent factors generated in MOFA. Bar colors represent the input dataset (ATAC-seq = coral, methylation = green, media-only RNA-seq = turquoise, Mtb-stimulated RNA-seq = blue, SNP = fuschia). (B) Sample weights on the first four MOFA factors were tested for RSTR vs LTBI using a linear mixed effects model corrected for age, sex and genetic kinship (FDR < 0.1). Black squares indicate groupwise means.

### Overlap analysis with additional latent variable integration methods

We next validated these MOFA findings using alternative data integration methods. Three additional data integrations were performed: two methods from the mixOmics package yielded two latent variables each, and the iClusterPlus package yielded three latent variables (Supplemental Figure 6). All four latent variables generated by the mixOmics methods and two generated in iClusterPlus significantly differed between LTBI and RSTR groups (FDR < 0.1, Supplemental Table 1). High-weighted features on each MOFA factor were selected proportional to the size and dataset contribution to total variance for each modality explaining >5% of variance on the factor. High-importance features on the latent factors generated in the non-MOFA methods were also compiled by selecting features with the top 10% absolute value on each significant latent variable, and features implicated in MOFA and at least two of the three additional methods were selected for functional enrichment (Table 3, Supplemental Tables 2 & 3). This overlap-based feature selection resulted in 111 unique HGNC gene symbols for Factor 1, 94 for Factor 2, 379 for Factor 3, and 307 for Factor 4 (Supplemental Table 4). These reduced lists represent features whose relationship to RSTR status is robust to data integration method.

**Table 3.** Summary of select high-weight MOFA features with high loadings on latent variables generated in at least two out of three alternative data integration methods.

| MOFA factor | Dataset (% variance explained on factor) | Number of select MOFA features | Number of features also identified in 2/3 non-MOFA methods (%) | Number of unique protein coding HGNC symbols |
| --- | --- | --- | --- | --- |
| Factor 1 | RNA Mtb (50%) | 177 | 41 (23%) | 40 |
|  | RNA media (48%) | 169 | 84 (50%) | 83 |
|  | Total (98%) | 346 | 113 (32%) | 111 |
| Factor 2 | ATAC-seq (98%) | 605 | 132 (22%) | 94 |
| Factor 3 | Methylation (91%) | 1198 | 372 (31%) | 345 |
|  | SNP (6%) | 94 | 77 (82%) | 35 |
|  | Total (97%) | 1292 | 449 (35%) | 379 |
| Factor 4 | SNP (59%) | 973 | 334 (34%) | 173 |
|  | Methylation (19%) | 249 | 114 (46%) | 106 |
|  | RNA Mtb (11%) | 40 | 22 (55%) | 20 |
|  | ATAC-seq (7%) | 41 | 13 (32%) | 11 |
|  | Total (96%) | 1303 | 483 (37%) | 307 |

### Agreement with individual analysis of integrated datasets

To assess the extent of agreement between the individual analyses of these data and the high-importance features identified with MOFA, the reduced feature lists were compared with the individual analyses dataset results (Figure 3, Supplemental Table 5). Two genes in the reduced Factor 1 MOFA feature list (NLRP3, IFNG) and five genes in the reduced Factor 4 feature list (FCAR, IRF1, IRF8, MXD1, SECTM1) were also among the genes found to be significant for the interaction between Mtb stimulation and LTBI/RSTR status in the previous RNA-seq only analysis(13). For Factor 4, five methylation probes in the MOFA reduced feature list were also implicated in the individual analyses of the methylation dataset. One of these probes was annotated to a region with 13 overlapping protocadherin genes, and another was annotated to two nearest genes (AC006077.3 and PCBD2). The other three were annotated to BRDT, ABLIM1, and PKD1L2. Overall, these results suggest that some lead findings from individual datasets were retained in the MOFA analysis, but that we are also able to identify novel signatures through the integration.

**Figure 3.**
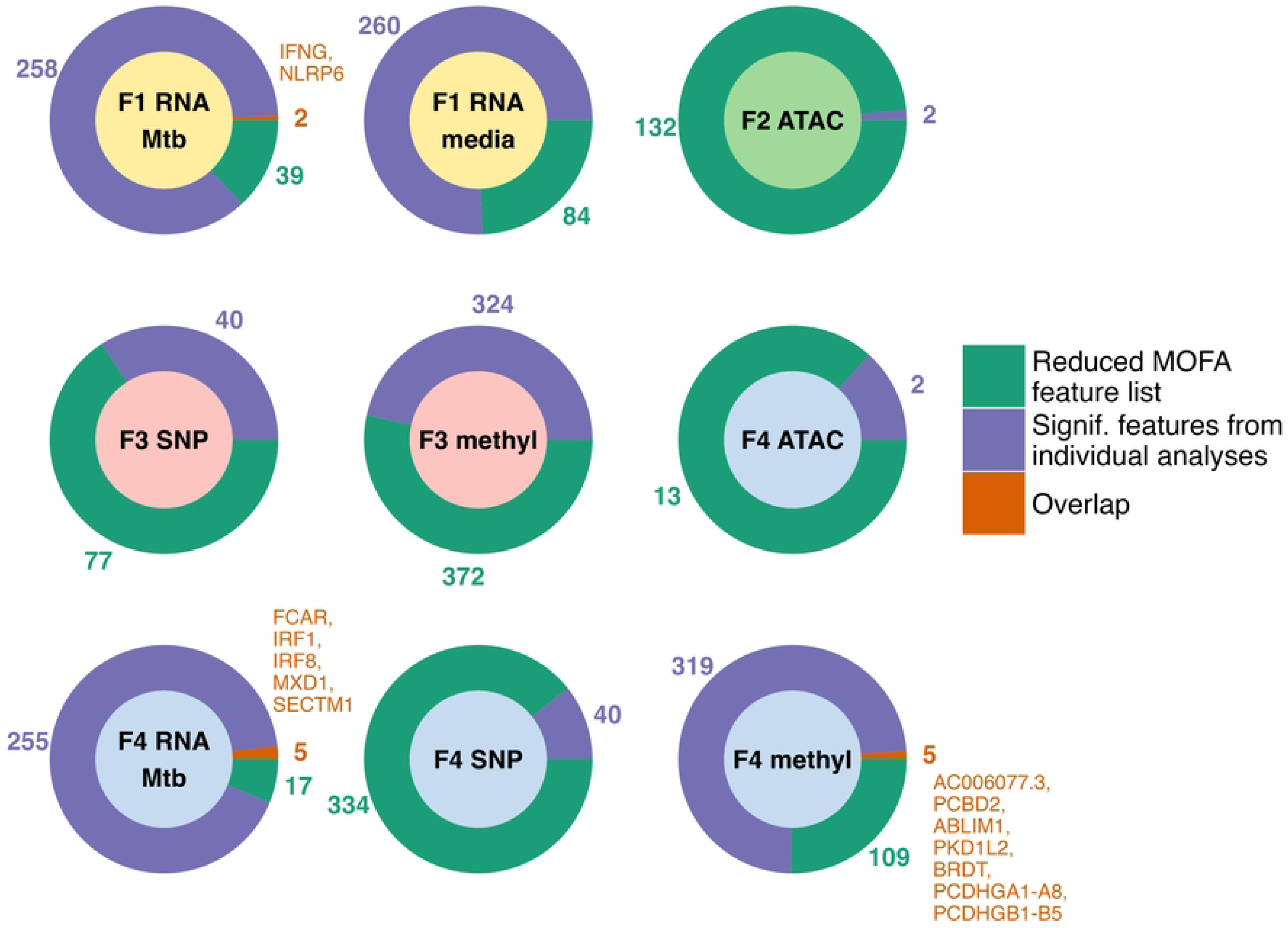
Features shared between MOFA reduced feature lists and results of individual analyses. Common features between reduced MOFA feature lists for Factors 1-4 (green) and significant features from individual analyses of the integrated datasets (purple; SNP: *P* < 5e-5; methylation: FDR < 0.2 in either probe list or list of probes in differentially methylated regions; ATAC-seq & RNA-seq: FDR < 0.2). Where there are overlapping features (orange), the unique HGNC symbols for the features are provided. In the case of methylation probes, some probes were annotated to more than one nearest gene. Inner circle color indicates the MOFA factor.

### MOFA factors are enriched for numerous and diverse functional pathways

We used pathway analysis to assess whether features within the MOFA factors were associated with biologic processes. Hypergeometric enrichment revealed hundreds of significantly enriched gene sets across the four factors, with many sharing common themes. There were 597 significantly enriched gene sets for Factor 1, 44 for Factor 2, 180 for Factor 3, and 183 for Factor 4 across Hallmark, canonical pathway, and gene ontology databases (FDR < 0.2, Supplemental Table 6, Supplemental Figure 7).

Significantly enriched gene sets were subjected to hierarchical clustering using the overlap coefficient to generate clusters of gene sets with similar gene membership (FDR < 0.2, Supplemental Table 7) and compared to generated in GO terms using the commonly employed semantic similarity method (Supplemental Figure 8, used by the rrvgo package(35)). The clusters generated on the overlap coefficient were thematically very similar to those generated on semantic similarity.

The media-only and Mtb-stimulated RNA-seq datasets contributed roughly equally to the list of features used in the Factor 1 enrichment. The largest cluster in the Factor 1 enrichment, Cluster F1-6, consisted of 136 gene sets primarily related to immune function (Supplemental Table 6). This cluster included the GO interferon gamma production pathway, which relates to the clinical definition of LTBI vs. RSTR. The six genes overlapping between this gene set and the MOFA reduced feature list (CD2, CD3E, CD96, GATA3, KLRK1, and NLRP6) were contributed by ten unique features from the media-only and Mtb-stimulated RNA seq datasets. In all cases, and in the case of IFN-γ itself, expression of these genes was higher in the LTBI subjects relative to the RSTR subjects (Figure 4). Cluster F1-1 (36 gene sets) also related primarily to immune function, particularly to T-cell activation and inflammatory responses. The largest clusters in Factors 2 and 3 included those related to cell-cell adhesion (F2-2, F2-4, F3-1) and cytoskeletal processes (F2-2, F3-2). Clusters containing at least 5 gene sets from the enrichment performed on the Factor 4 feature list are summarized in Table 4 and the member gene sets for the four largest clusters are displayed in Figure 5. The largest cluster, Cluster F4-4 (21 gene sets), as well as Clusters F4-30 (9 gene sets) and F4-20 (8 gene sets) contained gene sets with functions related to signal transduction, particularly G-protein signaling. Clusters F4-26 (14 gene sets) and F4-5 (9 gene sets) contained pathways related to cell-cell adhesion. Cluster F4-7 related to cell morphogenesis, and Cluster F4-8 related to structure- and tissue-level developmental pathways (both 13 gene sets). A handful of genes were highly prevalent (in > 50% of sets) within more than one gene set cluster. SRC had > 50% prevalence in five of these largest clusters. Three genes were prevalent in three of the summarized clusters (VAV2, GNA12, EPHB2) and four were in at least two clusters (HCK, BLK, DSCAM, PRKCZ) (Table 5). Taken together, these results suggest that MOFA Factors 2-4 are representing somewhat overlapping biological functions, primarily related to cell-cell adhesion, cell shape, and development of multicellular structures.

**Figure 4.**
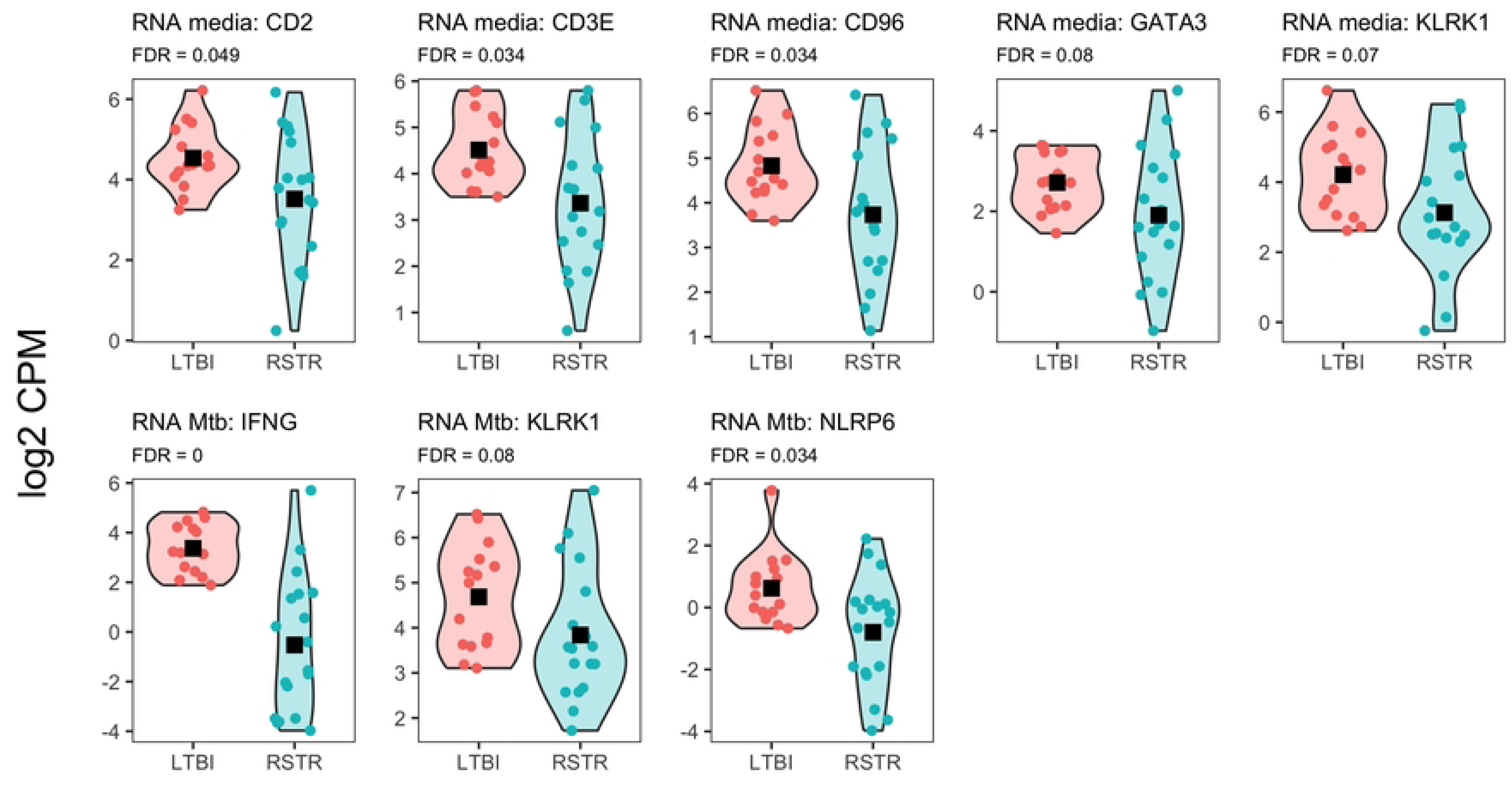
Expression of IFNγ-related genes is greater in LTBI than RSTR. Features in the Factor 1 reduced MOFA feature list belonging to the GO IFNγ production pathway are shown, in addition to IFNγ itself. IFNγ and NLRP6 are represented in the reduced feature list as part of the the Mtb-stimulated RNAseq dataset. KLRK1 is contributed by the both the Mtb-stimulated and media-only RNA-seq datasets, and CD2, CD3E, CD96, and GATA3 are contributed by the media-only RNA-seq dataset. For all features in both datasets, expression in greater in LTBI than RSTR (FDR < 0.1). Group differences tested by ANOVA; black squares represent groupwise means.

**Figure 5.**
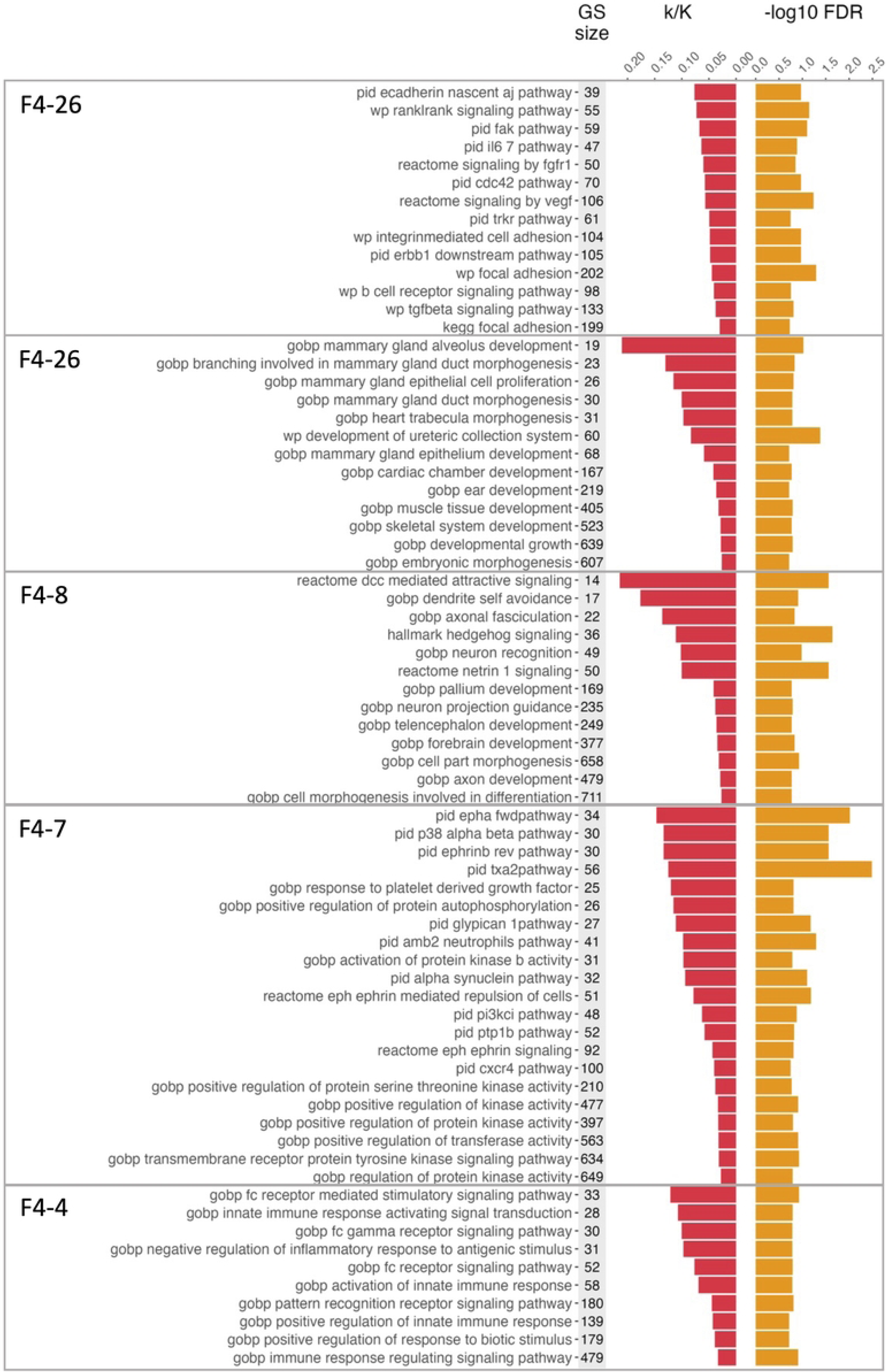
Significantly enriched gene sets in the largest clusters on Factor 4. Gene set names and sizes, and accompanying statistics for the four largest clusters of significantly enriched gene sets (FDR < 0.2) from hypergeometric testing of the Factor 4 reduced feature list. Colored boxes indicate cluster membership from the four largest gene set clusters generated via hierarchical clustering on the overlap coefficient.

**Table 4.** Summary of hypergeometric enrichment of select features on Factor 4 (FDR < 0.2, clusters with > 4 gene sets).

| Cluster | N gene sets | Cluster description | Genes in > 50% of gene sets | median k/K | median FDR |
| --- | --- | --- | --- | --- | --- |
| F4-4 | 21 | Signal transduction | SRC, BLK, HCK | 0.078 | 0.123 |
| F4-26 | 14 | Cell-cell adhesion & cell cycle | SRC, VAV2 | 0.053 | 0.107 |
| F4-7 | 13 | Cell morphogenesis | EPHB2, CNTN4, DSCAM | 0.041 | 0.148 |
| F4-8 | 13 | Structure & tissue-level development | BMP5, CHD7 | 0.059 | 0.165 |
| F4-3 | 10 | Innate immune response | HCK, SRC, FCN1 | 0.073 | 0.163 |
| F4-5 | 9 | Cell-cell adhesion | CDH18, CDH5, DSCAM, SLITRK1 | 0.039 | 0.162 |
| F4-30 | 9 | Signal transduction | GNA15, SRC, GNA12 | 0.067 | 0.079 |
| F4-20 | 8 | GTPase signaling, especially Ras-family signaling | ARHGEF18, ARHGEF3, VAV2, CDC42EP4, GNA12 | 0.050 | 0.140 |
| F4-13 | 7 | Epithelial cell differentiation | ALOX15B, AQP3, REG3G, ZBED2, HEY2, PALLD, PLEC, ST14 | 0.049 | 0.148 |
| F4-19 | 6 | Protein localization to plasma membrane | DPP10, EPHA3, EPHB2, SPTBN1, GBP1, NHLRC1, PRKCZ | 0.049 | 0.167 |
| F4-6 | 5 | Immune cell activation | PRKCZ, BCL6, IRF1, RSAD2, EBI3, PRKCQ | 0.044 | 0.194 |
| F4-12 | 5 | Smooth muscle contraction | GUCY1A1, GUCY1A2 | 0.056 | 0.151 |
| F4-14 | 5 | Hemostasis | SRC, ADRA2B, BLK, PRKCQ, TXK, VAV2, EPHB2, GNA12, JMJD1C, ST3GAL4 | 0.044 | 0.156 |
| F4-15 | 5 | Synaptic signaling | SYT10 | 0.054 | 0.169 |

**Table 5.** Genes with high prevalence across more than one large gene set cluster in Factor 4.

| Gene | Clusters | Dataset | Feature ID | MOFA Weight | Groupwise Statistics<sup>A</sup> |  | P-value |
| --- | --- | --- | --- | --- | --- | --- | --- |
|  |  |  |  |  | LTBI | RSTR |  |
| SRC | F4-4, F4-26, F4-3, F4-30, F4-14 | SNP | rs12329503 | 0.529 | 0: 12<br>1: 3<br>2: 0 | 0: 4<br>1: 11<br>2: 3 | 0.003 |
|  |  |  | rs6018088 | 0.493 | 0: 10<br>1: 5<br>2: 0 | 0: 4<br>1: 8<br>2: 6 | 0.011 |
|  |  |  | rs6018148 | 0.465 | 0: 10<br>1: 5<br>2: 0 | 0: 4<br>1: 10<br>2: 4 | 0.018 |
|  |  |  | rs6018257 | 0.662 | 0: 12<br>1: 1<br>2: 1 | 0: 4<br>1: 10<br>2: 4 | 0.002 |
| HCK | F4-4, F4-3 | SNP | rs4561724 | 0.459 | 0: 9<br>1: 6<br>2: 0 | 0: 3<br>1: 13<br>2: 2 | 0.025 |
| BLK | F4-4, F4-14 | SNP | rs2248932 | 0.558 | 0: 7<br>1: 8<br>2: 0 | 0: 4<br>1: 8<br>2: 6 | 0.037 |
| DSCAM | F4-7, F4-5 | SNP | rs1012854 | 0.964 | 0: 8<br>1: 7<br>2: 0 | 0: 1<br>1: 7<br>2: 10 | < 0.001 |
|  |  |  | rs11700509 | 0.677 | 0: 8<br>1: 4<br>2: 3 | 0: 2<br>1: 10<br>2: 6 | 0.031 |
| PRKCZ | F4-19, F4-6 | SNP | rs2803310 | 0.597 | 0: 10<br>1: 5<br>2: 0 | 0: 2<br>1: 10<br>2: 6 | 0.002 |
| VAV2 | F4-26, F4-20, F4-14 | Methylation | cg21223341 | -0.293 | 4.015 ± 0.122 | 3.554 ± 0.154 | 0.029 |
| EPHB2 | F4-7, F4-19, F4-14 | Methylation | cg13102231 | -0.237 | 2.759 ± 0.101 | 2.767 ± 0.170 | 0.970 |
| GNA12 | F4-30, F4-20, F4-14 | ATAC-seq | ID_chr7_2742848_2743374 | -0.112 | 2.152 ± 0.236 | 0.834 ± 0.233 | < 0.001 |

<sup>A</sup>Groupwise summary statistics are calculated as count by genotype for SNP data and means ± SE for all other data types. P-values represent chi-squared tests for SNP features and ANOVAs for other data types.

## Discussion

Mechanisms of resistance to *Mycobacterium tuberculosis* infection are not well understood. MOFA integration of SNP, methylation, chromatin accessibility, and transcriptomic datasets derived from a Uganda resister cohort reveals four nascent factors that differentiate subjects based on TST/IGRA status following TB exposure. High-importance features on Factor 1 were primarily enriched for pathways related to immune function, particularly inflammation, T cell responses, and interferon gamma responses. Factor 4 nearly perfectly discriminates RSTR from LTBI subjects and has meaningful contribution from four of the five integrated datasets. This factor was enriched for several pathways related to cell-cell adhesion, cell morphogenesis, and development of multicellular structures. Enrichments on Factors 2 and 3 showed similar themes, with top pathways relating to cell-cell adhesion and cytoskeletal processes. With this integrated dataset, the rigorous selection of important features through multi-integration overlap, and the functional enrichments performed on those features, our study provides a resource for hypothesis generation and a point of comparison for future investigations on the molecular mechanisms of Mtb resistance.

The two largest gene set clusters on Factor 1 were related to immune function, including pathways related to interferon gamma production. The Factor 1 MOFA reduced feature list was enriched for pathways including interferon gamma production, adaptive immune cell surface receptor production(36–39), regulation of NK cell surface receptors(39) involved in creating the immune synapse, and T cell differentiation(40–42). Expression of genes within these pathways was higher in LTBI relative to RSTR subjects in either or both of the RNA-seq datasets. Given that Factor 1 is weighted for the two transcriptomic datasets, it is likely this factor captures groupwise differences in expression of immune pathways in response to Mtb, particularly expression of adhesion molecules on adaptive immune cells. The definition of the RSTR phenotype is lack of TST/IGRA conversion following Mtb exposure, so this factor probably describes sources of variance in the canonical response that defines the clinical phenotype.

Factors 2, 3, and 4 contained functional enrichment of pathways related to cell adhesion, multicellular structures, and signaling. One possible interpretation of these results points to a relationship between Mtb resistance and cell-cell interactions such as in the early stages of the formation of the granuloma, a multicellular structure created through the aggregation and adhesion of immune cells which surround Mtb. This complex and dynamic structure is a hallmark pathologic feature of TB and represents the interface of host-pathogen interactions that define the outcome of host protection or progression of infection (43–46). Crucial to the early formation of this structure is the implementation of an epithelialization program involving recruitment of macrophages, changes in cell shape, and cell-cell adhesion(43). Additionally, a number of genes with a known role in early granuloma formation appear in the Factor 4 reduced MOFA feature list. SLC11A1 has been identified as one of seven genes with increased expression in TB granulomas relative to those formed in sarcoidosis, a non-infectious granulomatous disease(44). This gene encodes a divalent cation transporter involved in macrophage activation and has been implicated in TB pathogenesis in mouse and human studies(44,47–49). The highest weighted feature on the Factor 4 reduced feature list was a SNP annotated to DSCAM. This gene encodes a cell adhesion molecule expressed in the fetal brain. While the mechanism is not known, this gene has been previously associated with TB susceptibility(50). Several other SNPs in the reduced feature list for this factor were annotated to the tyrosine kinase-coding gene SRC. SRC and related tyrosine kinases (including BLK, HCK) have been investigated as possible drug targets for TB treatment(51) and have been specifically implicated in the regulation of granuloma formation(52). VAV2 encodes a guanine nucleotide exchange factor involved in cytoskeletal rearrangement. Granulomas are classically observed at later stages of disease pathogenesis when biopsy samples are obtained from individuals presenting with signs or symptoms of TB. Histopathology data from those who resist TST/IGRA conversion are not available to assess whether there are granuloma-equivalents or other types of multicellular structures. Taken together, these results point to possible differences in multicellular structures, early granuloma formation, or recruitment of cells to inflammatory foci in differentiating RSTR and LTBI subjects.

Biologic interpretation of these MOFA feature lists proved challenging due to the high number of enriched pathways. Hypergeometric enrichment of the reduced MOFA feature lists resulted in hundreds of significantly overrepresented gene sets across the four factors. To summarize, identify, and interpret major themes from these results, we developed madRich: a method for cross-database gene set clustering and annotation using hierarchical clustering on the overlap coefficient(34). Other packages exist for clustering output from pathway analyses, but these often rely on network-based methods which may be slower, more computationally demanding to implement, and more difficult to interpret(53,54). For all methods, an underlying distance metric is required to generate a useful clustering result. Some methods make use of sematic similarity as a distance metric, which performs very well in hierarchically-structured reference databases as seen in Gene Ontology gene sets(35,55). However, this metric cannot be used to summarize enrichment results from non-GO reference databases or results concatenated from enrichments against more than one database. Another commonly used metric for sparse binary clustering is Jaccard similarity(54,56), but this metric performs poorly on gene set data, because it punishes highly disparate set sizes, even if the smaller set is entirely nested within the larger. Instead, we utilized the overlap coefficient in our clustering algorithm. The overlap coefficient is the proportion of shared elements between two sets divided by the size of the smaller set. Clustering on this coefficient will result in better grouping of child gene sets with parents relative to the Jaccard coefficient in the case of hierarchical databases and allows compositionally similar gene sets to be grouped across databases. For our largest gene set clusters on Factor 4, madRich clustering of GO terms overlapped largely with rrvgo clustering, with madRich combining some rrvgo clusters (like rrvgo “wound healing” and “regulation of body fluid levels” combined into madRich “hemostasis”) and splitting others (like rrvgo “ear development” and “heart trabercula development” being split between madRich “structure and tissue development” and “cell morphogenesis”). But largely, the same themes emerged from the study of both sets of gene set clusters from the Factor 4 enrichment result. One advantage of rrvgo is the automation of cluster annotation. But as with the rrvgo cluster specifically annotated to “ear development,” which contains several multicellular structural morphogenesis pathways not exclusive to the ear, manual curation of these annotations is often necessary. Provided that some prefiltering is done to remove very large, broad gene sets, hierarchical clustering on the overlap coefficient is an effective way to summarize a complex enrichment result and glean relevant biological insights and has advantages over rrvgo, a commonly used alternative.

There are several limitations to the current work. First, the small sample size of 33 individuals is small and limits power. Although MOFA can interpolate missing datasets, these results are highly skewed when entire datasets are missing as opposed to individual features within an otherwise complete dataset. Additionally, the other integration methods used in the selection of top features do not allow for the interpolation of entire missing datasets for a subject. For these reasons, we decided to focus on the subset of subjects with complete data across the five integrated input datasets. Future work could investigate the extent to which these findings are generalizable to the full Uganda resister cohort or other TB cohorts. Second, when selecting top features for comparison across integration methods and downstream functional enrichment, cutoffs are imposed that are necessarily arbitrary. We selected cutoffs to include enough features to have an interpretable enrichment across all four significant factors and to have a reasonable contribution of features from the smaller RNA-seq and ATAC-seq datasets in comparison to the larger methylation and genetic datasets. We mitigated the arbitrary nature of this feature selection by using generous statistical cutoffs for the MOFA feature lists coupled with assessment with multiple integration methods. Finally, because of the different underlying data structures, the biologic directionality of relationships between clinical groups and functionally enriched gene sets are difficult to ascertain.

In summary, multi-omic factor analysis identified four latent variables with significant relationships to RSTR status. Feature lists derived from these variables showed functional enrichment for hundreds of gene sets across commonly used gene set databases including insights not derived from the individual datasets. We also provided a method of summarizing, visualizing, and annotating complex, cross-database functional enrichment results. These results can serve as a resource for hypothesis generation and cross-validation in future studies of the molecular mechanisms underlying resistance to Mtb infection.

## Data Availability

Preprocessing and analysis code will be made available in GitHub upon publication at https://github.com/hawn-lab/Uganda_RSTR_integration. Raw data are available through the NCBI database of Genotypes and Phenotypes (dbGaP) Data Browser (https://www.ncbi.nlm.nih.gov/gap/) under accession phs002445.v3.p1 but first must be approved by the data access committee (DAC) for the study site (see Supplemental Methods in [13])

## Acknowledgements

We thank the individual study participants of the Kawempe Community Health Study, study coordinators and the clinical and research staff including LaShaunda Malone, Keith Chervenak, Marla Manning, Dr. Mary Nsereko, Dr. Moses Joloba, Hussein Kisingo, Sophie Nalukwago, Dorcas Lamunu, Deborah Nsamba, Annet Kawuma, Saidah Menya, Joan Nassuna, Joy Beseke, Michael Odie, Henry Kawoya, Shannon Pavsek, Dr. E. Chandler Church, Anna Duewiger and Dr. Bonnie Thiel.

## Funding

The research was supported by the Bill and Melinda Gates Foundation (grant OPP1151836 to T.R.H., W.H.B., C.M.S., and H.M.K.), the National Institutes of Health (grant R01AI124348 to W.H.B., T.R.H., C.M.S., and H.M.K.; grant U01AI115642 to W.H.B., T.R.H., C.M.S., and H.M.K.; grant K24AI137310 to T.R.H.; R33AI138272 (to TRH, WHB, HMK, CMS), NIH grants K08AI143926 and T32AI007044 (to JDS), grant U19AI162583 (to HMK, WHB, CMS, and TRH), contract no. 75N93019C00071 to T.R.H., W.H.B., C.M.S., and H.M.K.; and contract no. NO1AI70022 to W.H.B., T.R.H., C.M.S., and H.M.K.). The funders had no role in the experimental design or analysis.

## Notes

### Competing Interest Statement

The authors have declared no competing interest.

## References

1. Global tuberculosis report. Geneva: World Health Organization; 2023.

2. Stein CM, Nsereko M, Malone LL, Okware B, Kisingo H, Nalukwago S, et al. Long-term Stability of Resistance to Latent Mycobacterium tuberculosis Infection in Highly Exposed Tuberculosis Household Contacts in Kampala, Uganda. Clin Infect Dis Off Publ Infect Dis Soc Am. 2019 May 15;68(10):1705–12.

3. Medawar L, Tukiman HM, Mbayo G, Donkor S, Owolabi O, Sutherland JS. Analysis of cellular and soluble profiles in QuantiFERON nonconverters, converters, and reverters in the Gambia. Immun Inflamm Dis. 2019 Aug 20;7(4):260–70.

4. Stein CM, Zalwango S, Malone LL, Thiel B, Mupere E, Nsereko M, et al. Resistance and Susceptibility to Mycobacterium tuberculosis Infection and Disease in Tuberculosis Households in Kampala, Uganda. Am J Epidemiol. 2018 Jul;187(7):1477–89.

5. Verrall AJ, Alisjahbana B, Apriani L, Novianty N, Nurani AC, van Laarhoven A, et al. Early Clearance of Mycobacterium tuberculosis: The INFECT Case Contact Cohort Study in Indonesia. J Infect Dis. 2020 Mar 28;221(8):1351–60.

6. Weiner J, Domaszewska T, Donkor S, Kaufmann SHE, Hill PC, Sutherland JS. Changes in Transcript, Metabolite, and Antibody Reactivity During the Early Protective Immune Response in Humans to Mycobacterium tuberculosis Infection. Clin Infect Dis Off Publ Infect Dis Soc Am. 2020 Jun 24;71(1):30–40.

7. Kroon EE, Correa-Macedo W, Evans R, Seeger A, Engelbrecht L, Kriel JA, et al. Neutrophil extracellular trap formation and gene programs distinguish TST/IGRA sensitization outcomes among Mycobacterium tuberculosis exposed persons living with HIV. PLoS Genet. 2023 Aug;19(8):e1010888.

8. Vorkas CK, Wipperman MF, Li K, Bean J, Bhattarai SK, Adamow M, et al. Mucosal-associated invariant and **γδ** T cell subsets respond to initial *Mycobacterium tuberculosis* infection. JCI Insight [Internet]. 2018 Oct 4 [cited 2024 Jul 11];3(19). Available from: https://insight.jci.org/articles/view/121899

9. Guwatudde D, Zalwango S, Kamya MR, Debanne SM, Diaz MI, Okwera A, et al. Burden of tuberculosis in Kampala, Uganda. Bull World Health Organ. 2003 Jan 20;81(11):799–805.

10. Stein CM, Zalwango S, Malone LL, Won S, Mayanja-Kizza H, Mugerwa RD, et al. Genome Scan of M. tuberculosis Infection and Disease in Ugandans. PLOS ONE. 2008 Dec 31;3(12):e4094.

11. McHenry ML, Benchek P, Malone L, Nsereko M, Mayanja-Kizza H, Boom WH, et al. Resistance to TST/IGRA conversion in Uganda: Heritability and Genome-Wide Association Study. eBioMedicine [Internet]. 2021 Dec 1 [cited 2024 Jan 23];74. Available from: https://www.thelancet.com/journals/ebiom/article/PIIS2352-3964(21)00521-1/fulltext

12. Dill-McFarland KA, Simmons JD, Peterson GJ, Nguyen FK, Campo M, Benchek P, et al. Epigenetic programming of host lipid metabolism associates with resistance to TST/IGRA conversion after exposure to Mycobacterium tuberculosis. bioRxiv. 2024 Mar 4;2024.02.27.582348.

13. Simmons JD, Dill-McFarland KA, Stein CM, Van PT, Chihota V, Ntshiqa T, et al. Monocyte Transcriptional Responses to Mycobacterium tuberculosis Associate with Resistance to Tuberculin Skin Test and Interferon Gamma Release Assay Conversion. mSphere. 2022 Jun 29;7(3):e00159–22.

14. Simmons JD, Van PT, Stein CM, Chihota V, Ntshiqa T, Maenetje P, et al. Monocyte metabolic transcriptional programs associate with resistance to tuberculin skin test/interferon-γ release assay conversion. J Clin Invest. 2021 Jul 15;131(14):e140073.

15. Chen C, Wang J, Pan D, Wang X, Xu Y, Yan J, et al. Applications of multi-omics analysis in human diseases. MedComm. 2023;4(4):e315.

16. Santiago-Rodriguez TM, Hollister EB. Multi ‘omic data integration: A review of concepts, considerations, and approaches. Semin Perinatol. 2021 Oct 1;45(6):151456.

17. Argelaguet R, Arnol D, Bredikhin D, Deloro Y, Velten B, Marioni JC, et al. MOFA+: a statistical framework for comprehensive integration of multi-modal single-cell data. Genome Biol. 2020 May 11;21(1):111.

18. Alda-Catalinas C, Bredikhin D, Hernando-Herraez I, Santos F, Kubinyecz O, Eckersley-Maslin MA, et al. A Single-Cell Transcriptomics CRISPR-Activation Screen Identifies Epigenetic Regulators of the Zygotic Genome Activation Program. Cell Syst. 2020 Jul;11(1):25–41.e9.

19. Rodriguez L, Pekkarinen PT, Lakshmikanth T, Tan Z, Consiglio CR, Pou C, et al. Systems-Level Immunomonitoring from Acute to Recovery Phase of Severe COVID-19. Cell Rep Med. 2020 Aug;1(5):100078.

20. Chang CC, Chow CC, Tellier LC, Vattikuti S, Purcell SM, Lee JJ. Second-generation PLINK: rising to the challenge of larger and richer datasets. GigaScience. 2015 Dec 1;4(1):s13742–015-0047–8.

21. Zheng X, Levine D, Shen J, Gogarten SM, Laurie C, Weir BS. A high-performance computing toolset for relatedness and principal component analysis of SNP data. Bioinformatics. 2012 Dec;28(24):3326–8.

22. Gogarten SM, Sofer T, Chen H, Yu C, Brody JA, Thornton TA, et al. Genetic association testing using the GENESIS R/Bioconductor package. Bioinformatics. 2019 Jul 22;35(24):5346–8.

23. NCBI [Internet]. [cited 2024 Jan 29]. Homo sapiens genome assembly GRCh38. Available from: https://www.ncbi.nlm.nih.gov/data-hub/assembly/GCF_000001405.26/

24. R Core Team. R: A language and environment for statistical computing [Internet]. Vienna, Austria: R Foundation for Statistical Computing; Available from: https://www.R-project.org/

25. Dill-McFarland KA, Mitchell K, Batchu S, Segnitz RM, Benson B, Janczyk T, et al. Kimma: flexible linear mixed effects modeling with kinship covariance for RNA-seq data. Bioinforma Oxf Engl. 2023 May 4;39(5):btad279.

26. Liberzon A, Birger C, Thorvaldsdóttir H, Ghandi M, Mesirov JP, Tamayo P. The Molecular Signatures Database (MSigDB) hallmark gene set collection. Cell Syst. 2015 Dec 23;1(6):417–25.

27. Rohart F, Gautier B, Singh A, Cao KAL. mixOmics: An R package for ‘omics feature selection and multiple data integration. PLOS Comput Biol. 2017 Nov 3;13(11):e1005752.

28. Mo Q, Wang S, Seshan VE, Olshen AB, Schultz N, Sander C, et al. Pattern discovery and cancer gene identification in integrated cancer genomic data. Proc Natl Acad Sci U S A. 2013 Mar 12;110(11):4245–50.

29. Mo Q, Shen R, Guo C, Vannucci M, Chan KS, Hilsenbeck SG. A fully Bayesian latent variable model for integrative clustering analysis of multi-type omics data. Biostatistics. 2018 Jan 1;19(1):71–86.

30. Dill-McFarland KA, Benson B, Cox MS, Janczyk T. SEARchways [Internet]. 2024. (GitHub Repository). Available from: https://github.com/BIGslu/SEARchways

31. Ashburner M, Ball CA, Blake JA, Botstein D, Butler H, Cherry JM, et al. Gene Ontology: tool for the unification of biology. Nat Genet. 2000 May;25(1):25–9.

32. The Gene Ontology Consortium, Aleksander SA, Balhoff J, Carbon S, Cherry JM, Drabkin HJ, et al. The Gene Ontology knowledgebase in 2023. Genetics. 2023 May 2;224(1):iyad031.

33. M K V, K K. A Survey on Similarity Measures in Text Mining. Mach Learn Appl Int J. 2016 Mar 30;3:19–28.

34. Cox MS. madRich [Internet]. Available from: https://github.com/UWISDOM/madRich

35. Sayols S. rrvgo: a Bioconductor package for interpreting lists of Gene Ontology terms. MicroPublication Biol. 2023:10.17912/micropub.biology.000811.

36. Binder C, Cvetkovski F, Sellberg F, Berg S, Paternina Visbal H, Sachs DH, et al. CD2 Immunobiology. Front Immunol. 2020 Jun 9;11:1090.

37. Menon AP, Moreno B, Meraviglia-Crivelli D, Nonatelli F, Villanueva H, Barainka M, et al. Modulating T Cell Responses by Targeting CD3. Cancers. 2023 Feb 13;15(4):1189.

38. Chiang EY, de Almeida PE, de Almeida Nagata DE, Bowles KH, Du X, Chitre AS, et al. CD96 functions as a co-stimulatory receptor to enhance CD8+ T cell activation and effector responses. Eur J Immunol. 2020;50(6):891–902.

39. Wensveen FM, Jelenčić V, Polić B. NKG2D: A Master Regulator of Immune Cell Responsiveness. Front Immunol [Internet]. 2018 [cited 2024 Feb 10];9. Available from: https://www.frontiersin.org/journals/immunology/articles/10.3389/fimmu.2018.00441

40. Zheng D, Kern L, Elinav E. The NLRP6 inflammasome. Immunology. 2021 Mar;162(3):281–9.

41. Radulovic K, Ayata CK, Mak’Anyengo R, Lechner K, Wuggenig P, Kaya B, et al. NLRP6 Deficiency in CD4 T Cells Decreases T Cell Survival Associated with Increased Cell Death. J Immunol. 2019 Jul 15;203(2):544–56.

42. Ho IC, Tai TS, Pai SY. GATA3 and the T-cell lineage: essential functions before and after T-helper-2-cell differentiation. Nat Rev Immunol. 2009 Feb;9(2):125–35.

43. Cronan MR. In the Thick of It: Formation of the Tuberculous Granuloma and Its Effects on Host and Therapeutic Responses. Front Immunol. 2022 Mar 7;13:820134.

44. Elkington P, Polak ME, Reichmann MT, Leslie A. Understanding the tuberculosis granuloma: the matrix revolutions. Trends Mol Med. 2022 Feb;28(2):143–54.

45. Ramakrishnan L. Revisiting the role of the granuloma in tuberculosis. Nat Rev Immunol. 2012 May 1;12(5):352–67.

46. Russell DG, Cardona PJ, Kim MJ, Allain S, Altare F. Foamy macrophages and the progression of the human tuberculosis granuloma. Nat Immunol. 2009 Sep;10(9):943–8.

47. Bellamy R, Ruwende C, Corrah T, McAdam KPWJ, Whittle HC, Hill AVS. Variations in the NRAMP1 Gene and Susceptibility to Tuberculosis in West Africans. N Engl J Med. 1998 Mar 5;338(10):640–4.

48. Hsu YH, Chen CW, Sun HS, Jou R, Lee JJ, Su IJ. Association of NRAMP 1 Gene Polymorphism with Susceptibility to Tuberculosis in Taiwanese Aboriginals. J Formos Med Assoc. 2006 Jan 1;105(5):363–9.

49. Vidal SM, Malo D, Vogan K, Skamene E, Gros P. Natural resistance to infection with intracellular parasites: isolation of a candidate for Bcg. Cell. 1993 May 7;73(3):469–85.

50. Chimusa ER, Zaitlen N, Daya M, Möller M, van Helden PD, Mulder NJ, et al. Genome-wide association study of ancestry-specific TB risk in the South African Coloured population. Hum Mol Genet. 2014 Feb 1;23(3):796–809.

51. Chandra P, Rajmani RS, Verma G, Bhavesh NS, Kumar D. Targeting Drug-Sensitive and - Resistant Strains of Mycobacterium tuberculosis by Inhibition of Src Family Kinases Lowers Disease Burden and Pathology. mSphere. 2016 Apr 13;1(2):e00043–15.

52. Reichmann MT, Tezera LB, Vallejo AF, Vukmirovic M, Xiao R, Reynolds J, et al. Integrated transcriptomic analysis of human tuberculosis granulomas and a biomimetic model identifies therapeutic targets. J Clin Invest [Internet]. 2021 Aug 2 [cited 2024 Feb 2];131(15). Available from: https://www.jci.org/articles/view/148136#SEC3

53. Ewing E, Planell-Picola N, Jagodic M, Gomez-Cabrero D. GeneSetCluster: a tool for summarizing and integrating gene-set analysis results. BMC Bioinformatics. 2020 Oct 7;21(1):443.

54. Bhuva DD, Tan CW, Liu N, Whitfield HJ, Papachristos N, Lee S, et al. vissE: A versatile tool to identify and visualise higher-order molecular phenotypes from functional enrichment analysis [Internet]. bioRxiv; 2022 [cited 2024 Feb 2]. p. 2022.03.06.483195. Available from: https://www.biorxiv.org/content/10.1101/2022.03.06.483195v1

55. Gu Z, Hübschmann D. simplifyEnrichment: A Bioconductor Package for Clustering and Visualizing Functional Enrichment Results. Genomics Proteomics Bioinformatics. 2023 Feb 1;21(1):190–202.

56. Jaccard P. The Distribution of the Flora in the Alpine Zone.1. New Phytol. 1912;11(2):37–50.

